# Antiviral activity of N-ω-Chloroacetyl-L-Ornithine on in vitro replication of Chikungunya virus

**DOI:** 10.1101/745455

**Authors:** Lucero Luna-Rojas, Amanda M. Avila-Trejo, Verónica Alcántara-Farfán, Lorena I. Rodríguez-Páez, Marvin Omar Pastor-Alonso, J. Leopoldo Aguilar-Faisal

## Abstract

The infections causes by Chikungunya virus (CHIKV), family Togaviridae, genus Alfavirus, have become a health problem around the world, due to its widespread occurrence, the high morbidity rate and the absence of vaccines or antiviral treatments. We analyzed a competitive inhibitor of ornithine decarboxylase, enzyme key in the biosynthesis of Polyamines (PAs), N-ω-chloroacetyl-L-ornithine (NCAO), as a possible inhibitor of CHIKV replication because intracellular polyamines participate in the transcription and *in vitro* translation of CHIKV. NCAO does not have any cytotoxic effect on C6/36 cells even at a concentration of 1000μM at 72h after post-exposure but in Vero cells the cytotoxic effect was presented above 380μM at 48h post-exposure, which was considered when determining the inhibitory effect on viral replication. We demonstrate that NCAO inhibits the replication of CHIKV in Vero and C6/36 cells in a dose-dependent manner causing a decrease in the PFU/mL of at least 4-logarithm (p <0.01) in both cell lines. Viral yields were restored by the addition of exogenous polyamines, mainly putrescine. HPLC analyses it shown that NCAO decrease content of intracellular PAs even though in infected cells mainly decreased spermidine and spermine, so NCAO inhibits CHIKV replication, by depleting the intracellular polyamines in Vero and C6/36 cells, suggesting that this compound is a possible antiviral in CHIKV infections.

**Importance:** The infections caused by Chikungunya virus (CHIKV), genus Alfavirus, have become a worldwide health problem, due to its high morbidity rate and the absence of vaccines or antiviral treatments. It is known that CHIKV use intracellular poliamines during transcription and traduction process, so the depletion of intracellular putrescine, spermidine and spermine reduce the viral replication. N-ω-chloroacetyl-L-ornithine (NCAO) is known as a competitive inhibitor of ornithine decarboxylase, key enzyme in Polyamines biosynthesis, but no studies have proven its activity on the inhibition of polyamine-dependent viral replication. Here we demonstrate NCAO inhibits in vitro CHIKV replication, so we propose NCAO as an antiviral candidate.

## Introduction

Chikungunya is a re-emergent virus transmitted by the bite of female mosquitoes of the genus Aedes, belongs to the family Togaviridae, genus Alfavirus, has a single strand of RNA of positive polarity with a genome measuring around ∼ 11.8 kb (Parashar, Cherian, 2014). Currently CHIKV infections are a public health problem, since its appearance in 2005, the virus has spread to new areas, causing disease on a global scale generating a high potential risk; Despite the reported cases and economic losses that have been generated in the affected areas, there are currently no vaccines and the current therapy is based on administration of analgesics, antipyretics and anti-inflammatory agents to relieve symptoms (Silva, Dermody., 2017); However, these treatments are deficient due to the chronic complications that can be generated, whose duration can range from several days to years, making them incapacitating. On the other hand, it has been shown that several DNA and RNA viruses, including Chikungunya, require intracellular polyamines to be able to carry out their replication, these are small, positively charged molecules derived from arginine that are involved in multiple cellular processes such as cell proliferation (Pegg, 2009; Kang et al., 1994), DNA and RNA stabilization (Thomas et al., 1995; Kumar et al., 2009), cellular stress (Mitchell et al., 1998) cell signaling (Iyer, Delcour.,1997) among other functions in eukaryotic cells; also, however, the virus has demonstrated its involvement in various processes for virus replication. Studies conducted by Gibson et al. demonstrated that the herpes simplex virus contains significant amounts of two major PAs in the virion: spermine restricted to the nucleocapsid and spermidine present in the viral envelope, both important to neutralize the viral DNA loads and thus facilitate the compaction and encapsidation of the virus. Genome (Gibson, Roizman.,1971). It has been found that the continuous synthesis of PAs is an absolute requirement for some viruses such as Vaccinia virus and Cytomegalovirus, since the biosynthetic pathway of polyamines is interrupted by an inhibitor such as DFMO, viral performance is seriously affected (Frugier et al., 1994). On the other hand, RNA viruses employ similar mechanisms to neutralize nucleic acids; however, studies in Semliki Forest virus showed that PAs are not present in the viral capsid, but rather, PAs seem to promote RNA synthesis, when there is a depletion of PAs, there is a parallel marked reduction in the activity of viral RNA polymerase from cells infected with Semliki virus (Tuomi et al., 1982); likewise Mounce et al. have shown that depletion of PAs in cells infected with viruses Chikungunya and Dengue virus via treatment with DFMO, or DENSpm (diethylnorspermine), an analogue of PAs that induces the expression of SSAT, affects translation, one of the first steps in viral replication, since the depletion of PAs limits the expression of non-structural proteins including the viral polymerase and with it the replication (Mounce et al., 2016a, 2016b). Due to the above, the pathway of PA biosynthesis could be a target in the development of possible antivirals against Chikungunya, in fact several pharmaceutical products directed to the biosynthetic pathway of the PAs have been subjected to clinical trials as is the case of the DFMO; however, despite the ability of this inhibitor to decrease the levels of intracellular PAs in vitro, it has certain drawbacks in vivo due to the toxicity that it generates at high concentrations, which is why the search for possible antivirals against Chikungunya continue; therefore, in this work we studied whether N-ω-chloroacetyl-L-ornithine (NCAO) can inhibit the replication of this virus since it is a competitive inhibitor of ornithine decarboxylase (ODC). Rodríguez-Paéz et al. demonstrated that the inhibition properties of this compound causes a cytotoxic and antiproliferative effect in human cancer cell lines because it is inhibited, it has been seen that the ODC that is overexpressed in many cancer cells have high ODC expression, observing that the compound exerts cytotoxic and antiproliferative effects in cancer cells but almost no cytotoxic effect is minimal in non-cancer non-control cells (Vargas-Ramírez et al., 2016); therefore, being NCAO is a promising compound to be used as a possible antiviral in the infections generated by CHIKV.

## Material and methods

Cell culture. Vero cells were maintained in Dulbecco’s modified Eagle medium / F-12 nutrient mixture (DMEM-F12; Sigma-Aldrich) supplemented with fetal bovine serum and Penicillin-streptomycin, incubated at 37 ° C and 5% CO2. The C6 / 36 cells were maintained in minimal essential medium (MEM, Sigma-Aldrich) supplemented with fetal bovine serum and penicillin streptomycin and incubated at 34 ° C in the absence of CO2.

### Treatment with NCAO

The 2000 mM NCAO was diluted with DMEM-F12 medium or MEM without supplementation and used at different concentrations 10, 100, 200, 300, 400, 500 and 1000 μM in the cell viability assays. For treatment with NCAO, the cells were reseeded with fresh medium 2.5% fetal bovine serum overnight and then incubated 48 hrs with the compound, before infection with Chikungunya virus. For the restitution tests of polyamines, putrescine, spermidine and spermine (Sigma-Aldrich) were diluted with DMEM-F12 or MEM medium and added to 10 μM after 48 h with the NCAO in cells infected with CHIKV.

### Viability tests

Vero and C6 / 36 cells were cultured as described above using the above-mentioned NCAO concentrations, and the cell viability was determined at 24, 48 and 72 h by the MTT method diluted with 1X PBS at 540 nm on ELISA reader.

### Viral infection and titration

Chikungunya virus was derived from the third passage in C6 / 36 cells and viral stocks stored at -70 °C. Infection in Vero and C6 / 36 cells maintained in the presence of the NCAO and during the addition of exogenous polyamines as designated. For infection, the virus was diluted in DMEM-F12 or MEM medium, respectively, without serum for a multiplicity of infection of 0.1; 0.25; 0.5 and 1 (MOI). The viral inoculum was added to the cell monolayer and incubated 1 hour with periodic movements, the monolayer was washed with PBS pH 7.4 then the culture medium was added, the supernatants were collected at 24 h post-infection for viral titration and the quantification of putrescine, spermidine and spermine by HPLC. For the quantification of viral yield, 150,000 Vero / well cells were plated and cultured, the plate was incubated for 24 hrs at 37 ° C with 5% CO2 and serial dilutions were made with a factor of 10 of the virus until dilution 10-7 placing each dilution in the corresponding wells, the plate was incubated at 37 ° C with 5% CO2 homogenizing every 10 min for one hour, after incubation, 0.5% carboxymethylcellulose (CMC) was added (Sigma-Aldrich) diluted in culture medium and incubated at 37 ° C with 5% CO2 until the presence of lytic plates was observed, cells were fixed with 10% formaldehyde for one hour, then stained with crystal violet-ethanol at 0.1% for 15 min, the viral titer was determined by counting the wells that presented between 10-100 lytic plates, above the 10-3 dilution.

Quantification of intracellular polyamines by HPLC in the presence of infection with chikungunya virus. Cells C6 / 36 and Vero were pre-treated at different concentrations of NCAO for 24 hrs and then infected in the presence of NCAO at MOI of 0.5 and 1 with CHIKV, supernatants and cell debris were obtained at 24 hrs post-infection. For quantification by HPLC, the pre-treatment was performed with trichloroacetic acid (Sigma-Aldrich) 50% and 1,8 diaminooctane (Sigma-Aldrich), 1 mM. The benzoylation of the polyamine standards and the samples was carried out with NaOH (Sigma-Aldrich) 2 M and benzoyl chloride (Sigma-Aldrich) 50%, the extraction was carried out with chloroform-HPLC (Sigma-Aldrich) and the compounds were evaporated with a stream of nitrogen. The HPLC analysis was performed using a C18 reverse phase column, Agilent 5 μm, of 250 x 4.6 mm, the mobile phase was a mixture of: methanol-water (60:40), eluted with a linear gradient at a speed flow rate of 0.4 mL / min, detection was performed at 229 nm at room temperature in the Agilent 1260 equipment (Morgan, 1998; Arisan et al., 2002)

## Results

Cell viability assays by MTT performed on Vero and C6 / 36 cells at 24 h (Fig.1A and 1D), show that the compound does not exert cytotoxic effect even at a concentration of 1000 μM, at 48 and 72 hrs show that the compound does not exert cytotoxic effect even at a concentration of 1000 μM in C6 / 36 cells (Fig. 1E and 1F), however; In Vero cells, a decrease in cell viability was observed after 48 hrs at concentrations above 300 μM (ANOVA * p <0.05) (Fig.1B). The EC50 of NCAO was determined in Vero cells at 48 and 72 hrs obtaining values of 386.12 and 323.8 μM of NCAO respectively (Fig.2A and 2B).

**Figure 1.**
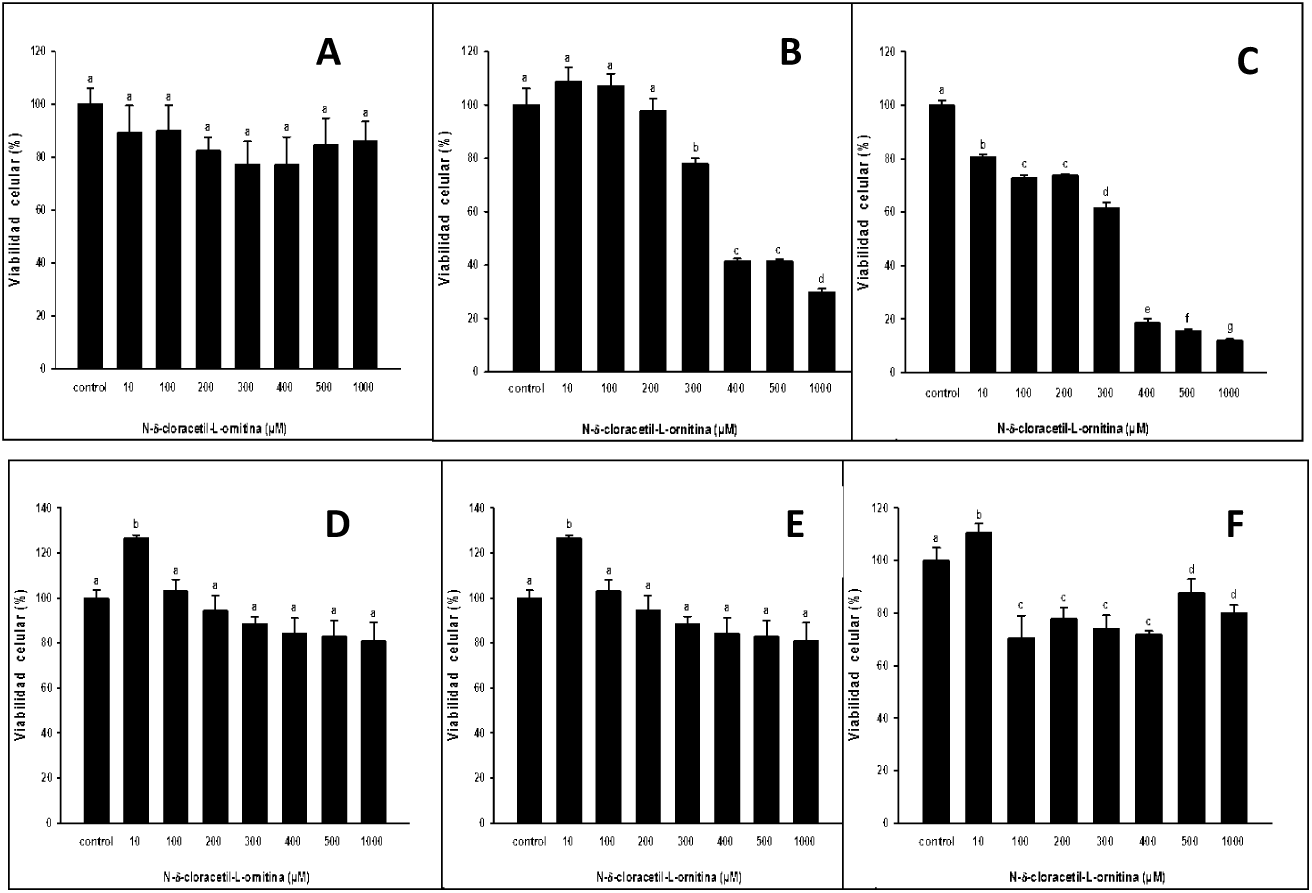
Viability of Vero and C6 / 36 cells treated with different concentrations of N-ω-chloroacetyl-L-ornithine. **A:**Cells were treated with NCAO growths at 24 B: 48 and C: 72 h using the MTT reduction method. D: C6 / 36 cells were treated with NCAO growths at 24, E: 48 and F: 72 h using the MTT reduction method (ANOVA P <0.05 n = 9). Different letters between groups.

**Figure 2.**
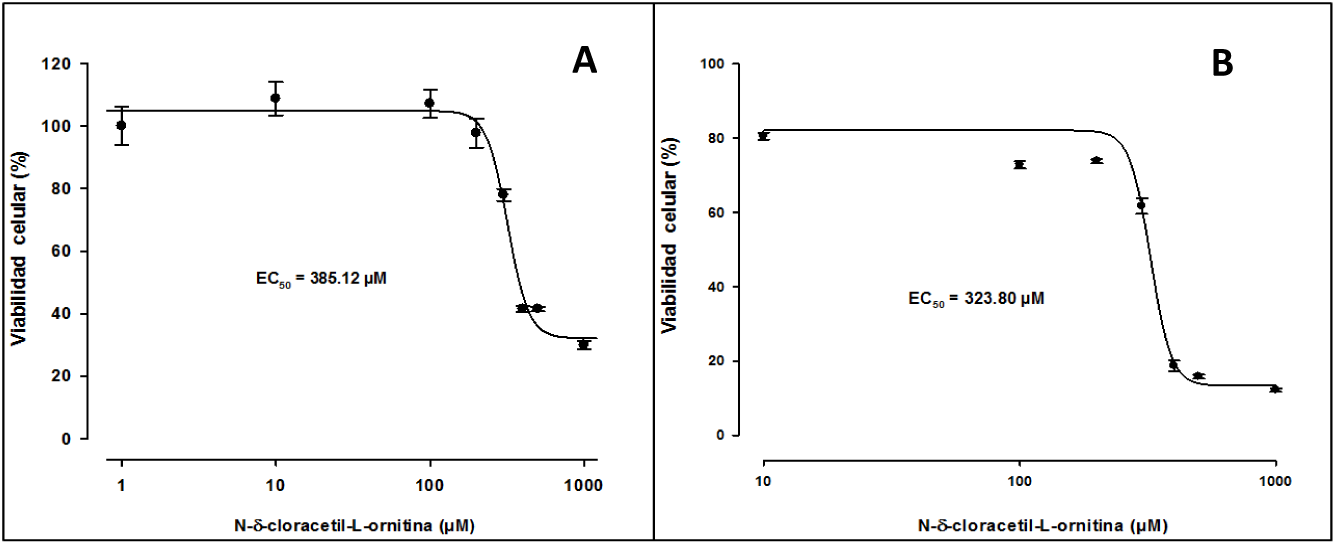
Concentration curves-EC_50_ response for Vero cells treated with NCAO. A: EC50 = 386.12 μM at 48h, and B: EC50 = 323.80 μM at 72 h.

Several studies have identified ODC as a key enzyme in the biosynthesis of polyamines, which are necessary for the replication of several viruses, including CHIKV (Mouce et al., 2016a, 2016b). To determine that the replication of the Chikungunya virus is affected by the depletion of the PAs in the presence of the NCAO, a competitive inhibitor of the ODC (Rodríguez-Páez et al., 1998; Correa-Basurto et al., 2009), pre-treatment of Vero and C6 / 36 cells was performed at two concentrations of NCAO 100 and 500 μM for 24 hrs before infection with CHIKV at MOI of 0.1; 0.25; 0.5 and 1, respectively (Fig. 3A and 3B). At 24 hrs post-infection (hpi), the PFU were determined and a significant reduction (ANOVA * p <0.01) was found in the viral titers in both concentrations in C6 / 36 cells compared to the control (Fig.3B). But nevertheless; in Vero cells only significant difference was found at a concentration of 500 μM (ANOVA *p <0.01) (Fig.3A).

**Figure 3.**
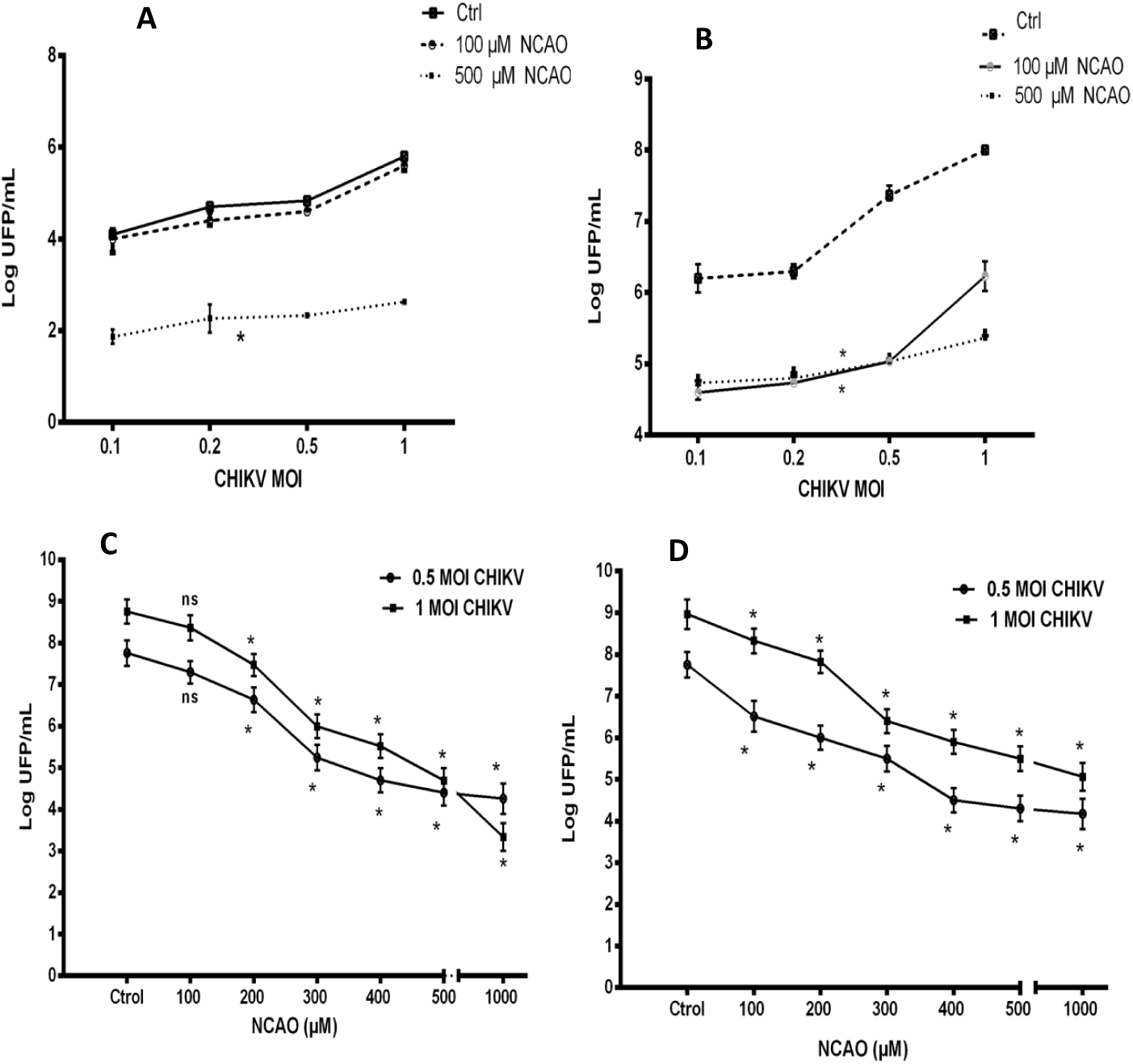
The NCAO affects the replication of Chikungunya virus. A and B: Vero and C6 / 36 cells pretreated at 100 and 500 μM of NCAO for 24 hrs and subsequently infected at 0.1; 0.25; 0.5; and 1 MOI with CHIKV, the titers were determined 24 hrs post-infection in Vero cells. C: Vero and D: C6 / 36 cells pretreated at 100, 200, 300, 400, 500 and 1000 μM of NCAO before and during infection with CHIKV at 0.5 and 1 MOI, titers were determined 24 hpi in Vero cells (ANOVA * p <0.01, n = 3, ns = No significant difference).

According to the previous results, it was considered to perform the kinetics of inhibition at the MOI of CHIKV 0.5 and 1 in Vero and C6 / 36 cells. In addition, it was considered to analyze the effects on the replication of the virus in the presence of NCAO even after pre-treatment, because it has been described that the enzyme presents a rapid turnover in mammalian cells, promoting the continuous synthesis of enzyme, which it would allow to avoid the effect of the compound (Pegg, 2009). The NCAO treatment was performed at the different concentrations 24 hrs before infection, and during the infection with the CHIKV, titers were measured at 24 hpi; a significant reduction was found in the titers from the 200 μM concentration in Vero cells (Fig.3C) and 100 μM in the C6 / 36 cells (Fig. 3D) observing a dose-dependent inhibition, since as the concentration of NCAO viral titers decreased (ANOVA * p <0.01) (Fig.3C and 3D).

To verify that the Chikungunya virus requires PAs to carry out its replication, exogenous polyamines putrescine, spermidine, spermine and the mixture thereof were added to the culture of Vero and C6 / 36 cells pre-treated with 300 μM NCAO and infected with the CHIKV at MOI 0.5 and 1.0; a restitution was observed in the viral titers in both cell lines when adding the exogenous PAs; the viral titers were similar to those obtained in the cells not treated with the compound, in comparison with the cells to which only the NCAO was added (ANOVA * p <0.01) (Fig.4A and 4B)

To confirm that CHIKV requires PAs for replication and that this is affected, due to NCAO treatment. The concentration of intracellular PAs in Vero and C6 / 36 cells without infection and infected with CHIKV was determined by HPLC (Fig.5). A decrease in the content of intracellular PAs in the presence of NCAO and during infection was observed at the two MOIs tested with respect to control in Vero and C6 / 36 cells (ANOVA * p <0.01) (Fig. 5A, 5B, 5C and 5D). In addition, it was observed that the content of intracellular PAs is higher in Vero cells than in C6 / 36 cells, and that the content of Putrescine decreases significantly in each of the conditions tested with respect to the control; however, it is the polyamine found in greater quantity, even in the presence of the NCAO. The content of spermidine and spermine decreased even in the presence of putrescine, the precursor PA. Interestingly, a significant decrease in spermidine content was observed during infection with CHIKV in both cell lines (ANOVA * p <0.01) (Fig. 5A, 5B, 5C and 5D).

**Figure 4.**
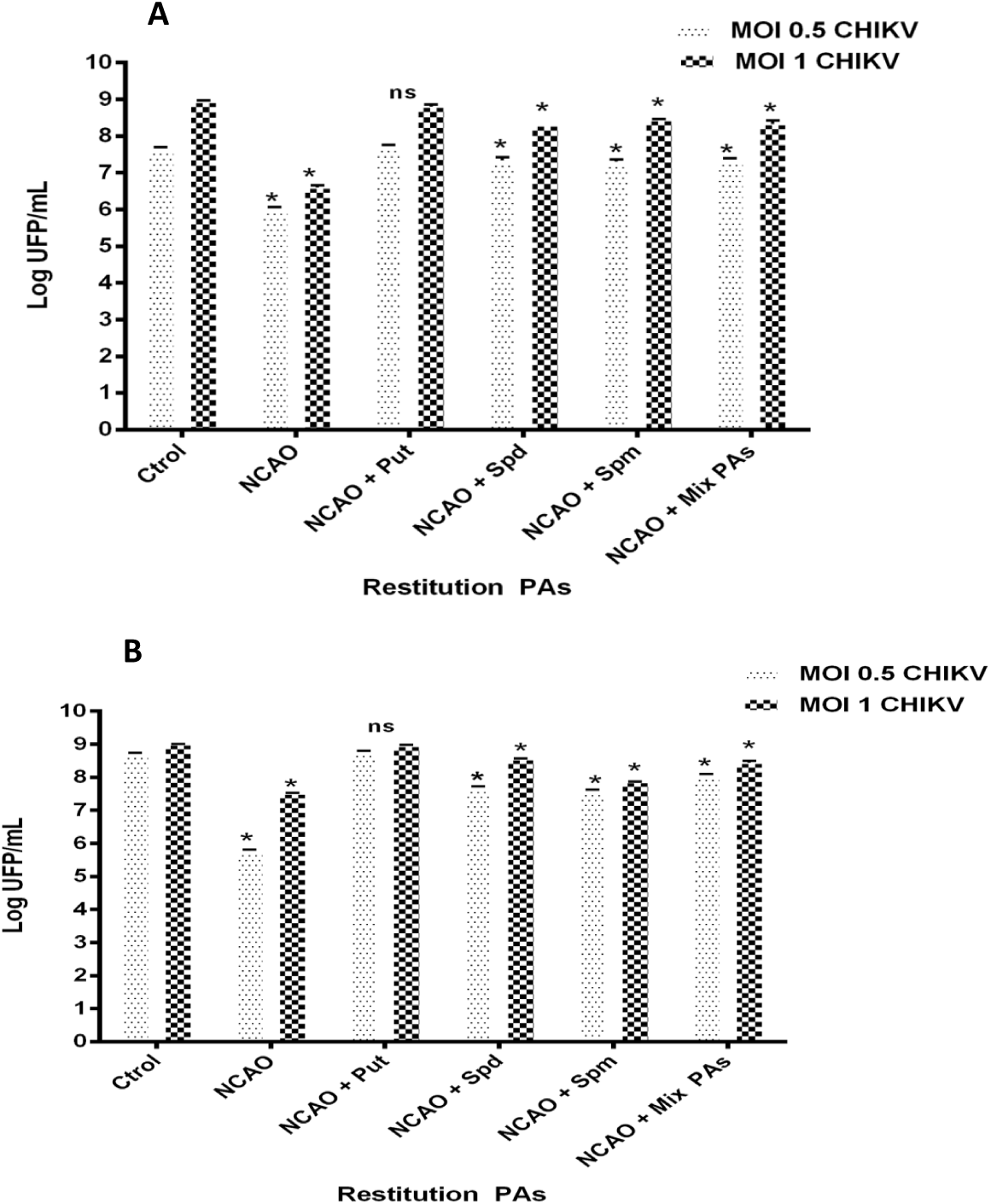
Restitution of CHIKV titers by addition of PAs in Vero and C6 / 36 cells. **A**:Vero and B: C6 / 36 cells pretreated at 300 μM of NCAO and addition of 10 μM PAs during infection with CHIKV at 0.5 and 1 MOI, titers were determined 24 hpi in Vero cells. A triplicate test was carried out (ANOVA * p <0.01 n = 3, ns = No significant difference).

**Figure 5.**
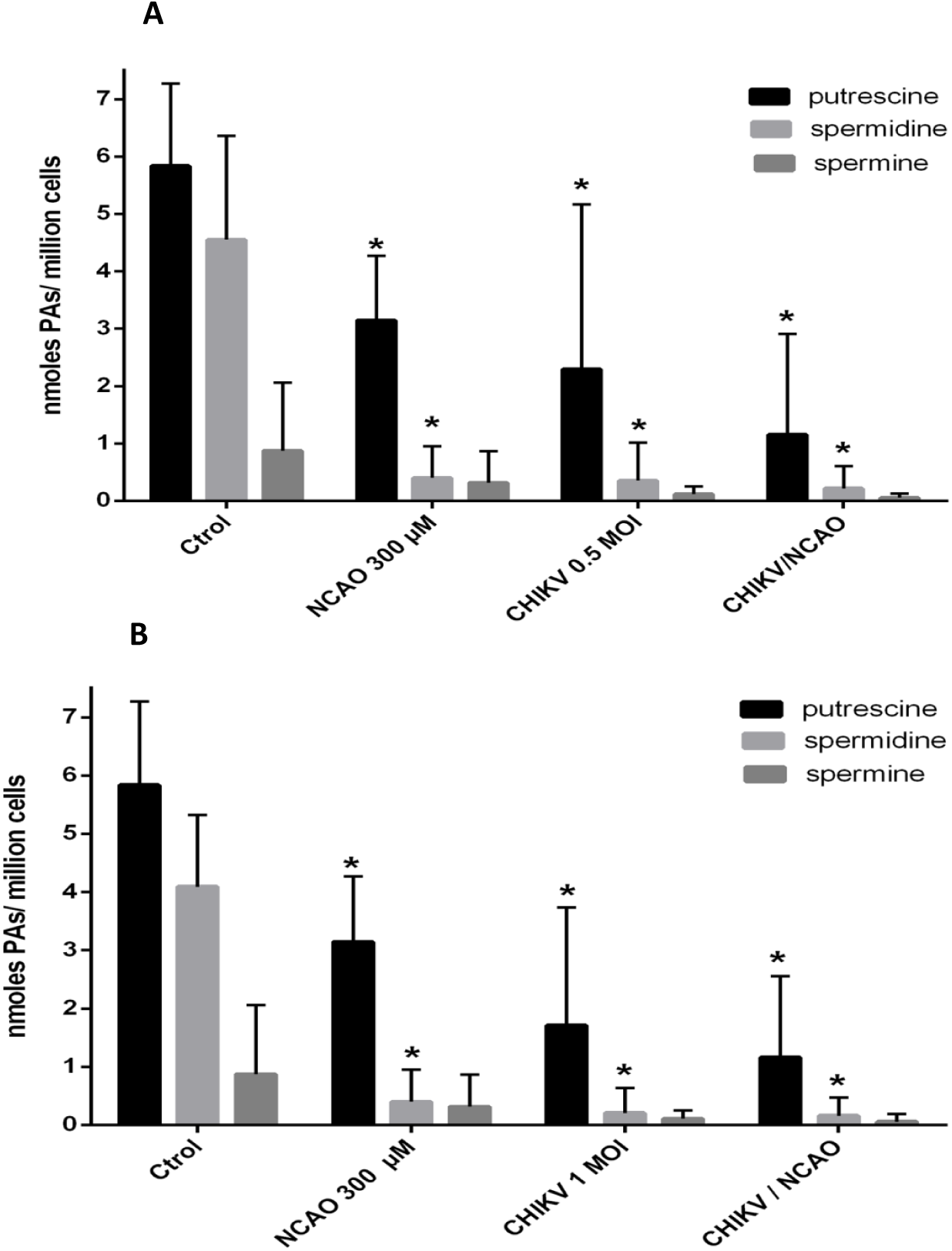
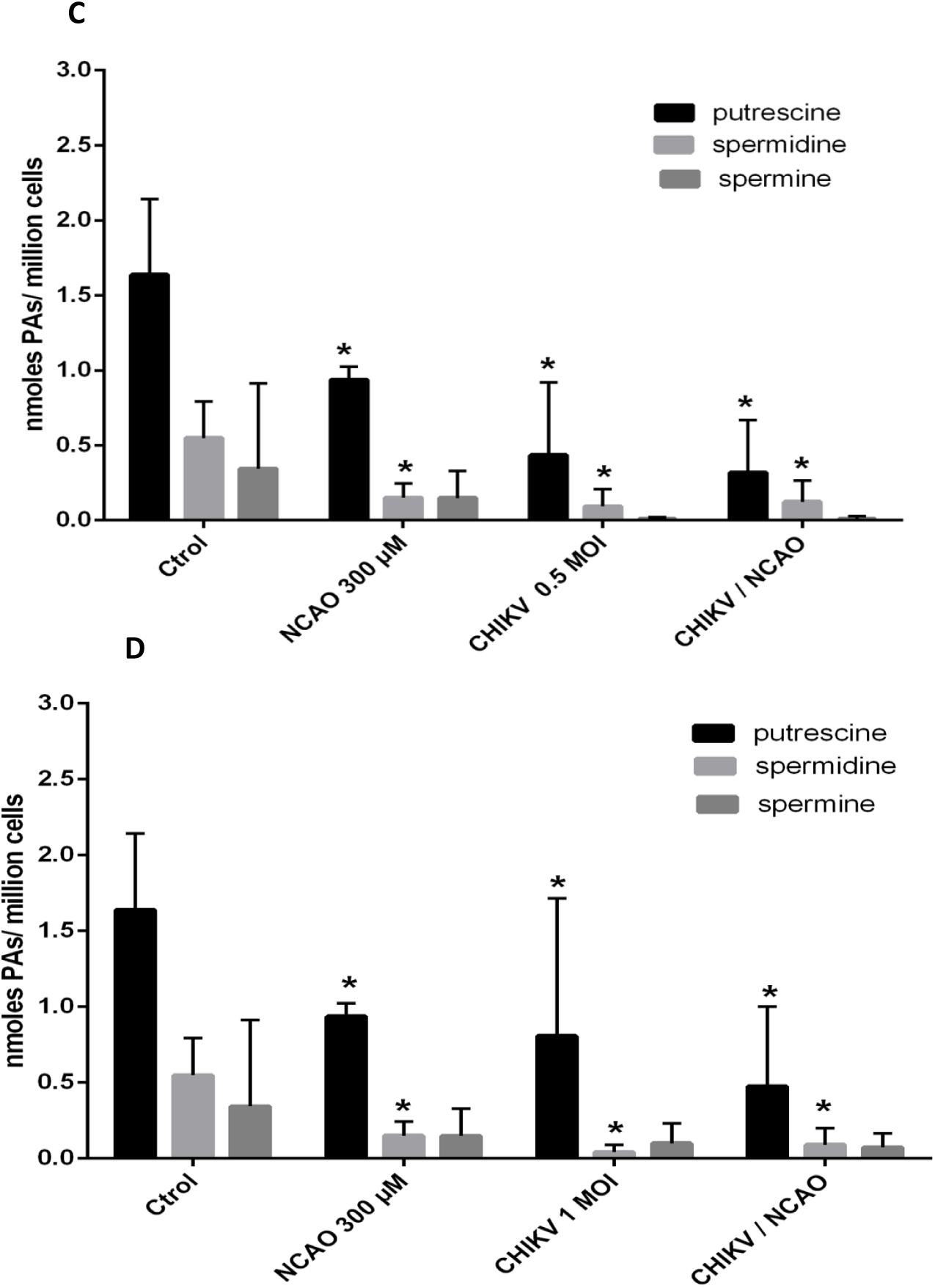
Content of intracellular PA in Vero and C6 / 36 cells. **A:**The Vero cells were previously treated with 300 μM of NCAO and infected to 0.5 and B: 1 MOI with CHIKV, the determination of intracellular PA content was done by HPLC. C: C6 / 36 cells pretreated to 300 μM of NCAO and infected at 0.5 and D: 1 MOI with CHIKV. The determination of the quantification of the content of the intracellular PA was performed by HPLC. (ANOVA * p <0.05 n = 3, ns = No significant difference).

## Discussion

CHIKV infections are a public health problem, due to the increase in the number of cases that has spread across several continents (Raiza et al., 2018). The widespread dissemination of CHIKV, the associated high morbidity rate, the lack of available treatments and the limited knowledge about the molecular mechanisms involved in the replication process open up an opportunity panorama for the investigation of these topics that allow the development of antivirals or effective vaccines for the control of the infection generated by CHIKV (Silva, Dermody., 2017). Several compounds have been proposed, including chloroquine (Brighton, 1984), ribavirin (Leyssen et al., 2008), arbidol (Delogu et al., 2017), harringtonine (Kaur et al., 2013), suramina (Albulescu et al., 2015) and silymarin (Lani et al., 2015) that act in different stages of the viral replication cycle, however, have certain disadvantages that make them ineffective for use in CHIKV infections. (Mishra et al., 2016). Mouce et al., 2016a, 2016b showed that PAs, putrescine, spermidine and spermine are important in the replication of a large variety of RNA viruses, including Chikungunya, as well as their participation in other molecular processes such as the packaging of the viral genome; in DNA viruses the polyamines stabilize the negatively charged genome within the virion particle, among them, the bacteriophage T7 virions cytomegalovirus (CMV) (Kalejta, 2008), and vaccinia virus are found (Lanzer, Holowezak., 1975). Therefore, this study evaluated whether N-ω-chloroacetyl-L-ornithine has an antiviral effect by decreasing or inhibiting the replication of CHIKV since it is a competitive inhibitor of Ornithine Decarboxylase, thereby decreasing the biosynthesis of PAs (Vargas-Ramírez et al., 2016). In this work it was demonstrated, through kinetic inhibition assays, that the NCAO inhibits the replication of the Chikungunya virus in Vero and C6 / 36 cells in a dose-dependent manner (Fig.3C and 3D) and that this inhibition is generated by depletion of the intracellular PAs in both cell lines as demonstrated by the quantification of PAs by HPLC (Fig. 5). In such a way, that there is a relationship between the content of PAs and viral replication. These results match with that described by Mouce et al., 2016a, 2016b, who demonstrated that there is a decrease in viral replication due to the depletion of polyamines in the cell lines C6 / 36, BHK21 and Vero cells induced by inhibitors of PA synthesis. the suicide inhibitor of ODC, difluoromethylornithine (DFMO), or diethyl-nospermine (DENSpm), an activator of SAT1, which induces the exhaustion of spermidine and spermine by the induction of SAT1 (Mounce et al., 2016a, 2016b); they also described that the decrease in PAs generated by these compounds exerts a negative effect in vivo and in vitro on the replication of various RNA viruses including Chikungunya (Mounce et al., 2017a, 2017b). However, the great potential of the DFMO to be used to combat viral infections would be overshadowed by adverse effects that would occur with the use of high doses of this compound to counteract the replacement of the ODC (FL Meyskens et al., 1986), and the administration of type I interferon to induce STAT1 activity could represent a high cost. In this sense, the NCAO being a less toxic compound could be considered in vivo studies as an antiviral in the infection with Chikungunya virus.

In the present work it was proved that CHIKV requires the polyamines for its replication when adding exogenous PAs in the cells pre-treated with NCAO, a restitution of the CHIKV titres was observed in both cell lines (Fig.4) by which, corroborated that the compound acts by inhibiting viral replication, by blocking the pathway of biosynthesis of PAs, this supports the findings obtained by Mouce et al., 2017a, 2017b where it shows the restitution of viral titers in various viruses, among them Chikungunya, when biogenic PAs are added, more than when the synthetic ones are added, after pretreatment with DFMO (Mounce et al., 2016a, 2016b). In order to know which of the PAs would have the best effect, putrescine, spermidine or spermine were individually added, finding better restitution of the viral titers with putrescine. This suggests that putrescine is more easily captured by the cells and used as a precursor for the synthesis of the other polyamines, which could be demonstrated by quantifying spermidine and spermine in the cells exposed to putrescine.

To rule out that viral replication was diminished by low viability, the viability of C6 / 36 cells in the presence of NCAO was determined and it was found that it does not exert cytotoxic effect during the treatment times (Fig. 1D, 1E and 1F); however, viability in Vero cells decreases (Fig. 1A, 1B and 1C) at concentrations above the EC50 at 48 and 72 h (Fig.2). Interestingly, the intracellular PAs are in greater quantity in the Vero cells than in the C6 / 36 cells (Fig.5), which opens questions regarding the efficiency of replication, since this virus requires PAs and is replicated more efficiently in C6 / 36 cells. Furthermore it has been described that the enzymes involved in the synthesis and catabolism of PAs and catabolism are conserved throughout the kingdoms; recent studies of polyamines in mosquitoes *Aedes aegypti* showed that it is probable that the content of PAs probably depends on the enzyme aaNAT5b an acetyltransferase (SAT); so it could act as a viral restriction factor, by decreasing intracellular PAs (Guan et al., 2018); However, there are likely to be other mechanisms in the cells that allow the virus to replicate successfully in the absence of polyamines.

Regarding the concentration of PAs in Vero and C6 / 36 cells during infection with CHIKV (Fig. 5), a decrease in the presence of NCAO was observed at the two MOIs tested (Fig. 5A, 5B, 5C and 5D).), which was expected since the NCAO decreases the pools of intracellular PAs, by inhibiting the ODC (Vargas-Ramirez et al., 2016); In addition, spermidine and spermine are diminished because the virus consumes these molecules during their replication; in several experimental studies it has been demonstrated that the activity of the RNA-dependent RNA polymerase (RdRP) of Chikungunya (Mounce, et al., 2017a) and Semliki Forest virus (SFV) (Tuomi et al., 1982) it is diminished in the absence of polyamines and its activity is restored by adding spermidine (Mounce, et al.,2017a, 2017b). Likewise, it is known that spermidine is important in one of the most critical steps of viral replication, translation, since the depletion of PAs limits the synthesis of non-structural proteins including viral polymerase (Mounce et al., 2016a, 2016b), since spermidine plays an important role specifically in the initiation and lengthening of translation, by hypofunctioning the 5A initiation factor (elF5A) (Nishimura et al., 2005). In this work it was observed that putrescine is found in greater quantity, even in the presence of the NCAO, which could suggest that the cell has other mechanisms to assure the content of this PA. Currently, the mechanism by which PA transport is carried out has not been well elucidated; however, it is known that the transport of putrescine to the interior of the cell could be through the exporter SLC3A2 present in the cell membrane (Nowotarski et al., 2013). Interestingly, in the treatment with NCAO the spermidine and spermine content decreases even in the presence of putrescine, the precursor PA, which suggests that NCAO could interact with some other enzyme of the biosynthetic route, in addition to the ODC.

## Conclusion

N-ω-chloroacetyl-L-ornithine (NCAO) inhibits the replication of the Chikungunya virus in Vero and C6 / 36 cells in a dose-dependent manner by depleting the three intracellular polyamines; although all three are important in the replication of this virus, spermidine may have a more complex role in the process of translation and transcription of the virus. Therefore, subsequent studies will provide more information about the possible use of this compound or some derivative as an antiviral.

## Ethical Approval

The project was reviewed and approved by the biosafety committee of the Higher School of Medicine-NIP.

## References

1. Albulescu IC, Van HM, Wolters LA., et al. Suramin inhibits chikungunya virus replication through multiple mechanisms. Antiviral Res. 2015;121, 39–46. https://doi.org/10.1016/j.antiviral.2015.06.013

2. Brighton, S. W. Chloroquine phosphate treatment of chronic Chikungunya arthritis. An open pilot study. S Afr Med J. 1984; 66(6):217–218.

3. Correa-Basurto J, Rodríguez-Páez L, Aguilar-Moreno ES., et al. Computational and experimental evaluation of ornithine derivatives as ornithine decarboxylase inhibitors. Medicinal Chemistry Research. 2009; 18(1):20–30. https://doi.org/10.1007/s00044-008-9103-6

4. Delogu I, Patorino B, Baronti C., et al. In vitro antiviral activity of arbidol against Chikungunya virus and characteristics of a selected resistant mutant. Antiviral Res. 2011; 90(3): 99–107. https://doi.org/10.1016/j.antiviral.2011.03.182

5. Frugier M, Florentz C, Hosseini MW., et al. Synthetic polyamines stimulate in vitro transcription by T7 RNA polymerase. Nucleic Acids Research. 1994;22(14): 2784–2790. https://doi.org/10.1093/nar/22.14.2784

6. Gibson W, Roizman B. Compartmentalization of spermine and spermidine in the herpes simplex virion. Proc Nactl Acad Sci USA. 1971;68(11):2818–2821. https://doi.org/10.1073/pnas.68.11.2818

7. Guan H, Wang M, Liao C., et al. Identification of aaNAT5b as a spermine N-acetyltransferase in the mosquito, Aedes aegypti. PLoS ONE. 2018;13(3): 1–13. https://doi.org/10.1371/journal.pone.0194499

8. Iyer R, Delcour AH. Complex inhibition of OmpF and OmpC bacterial porins by polyamines. J Biol Chem. 1997; 272(30):18595–18601. https://doi.org/10.1074/jbc.272.30.18595

9. Kaur P, Thiruchelvan M, Lee RC., et al. Inihibition of Chikungunya virus replication by harringtonine, a novel antiviral that supresses viral protein expression. Antimicrob Agents Chemother. 2013; 57(1):155–67 https://doi.org/10.1128/AAC.01467-12

10. Kalejta RF. Tegument Proteins of Human Cytomegalovirus. Microbiol Mol Biol Rev.,m 2008;72(2):249–265. https://doi.org/10.1128/JVI.01347-16

11. Kumar N, Basundra R, Maiti S. Elevated polyamines induce c-MYC overexpression by perturbing quadruplex-WC dúplex equilibrium. Nucleic Acids Res. 2009;37 (10): 3321–3331. https://doi.org/10.1093/nar/gkp196

12. Meyskens F., et al. A phase II study of alpha-difluoromethylornithine (DFMO) for the treatment of metastatic melanoma. Investigational New Drugs, 1986 4(3), 257–262.

13. Mitchell JL, Judd G, Leyser A., et al. Osmotic stress induces variation in cellular levels of ornithine decarboxylase-antizyme. Biochem. J. 1998;329(Pt 3): 453–459. http:// DOI: 10.1042/bj3290453

14. Mishra P, Kumar A, Mamidi P., et al. Inhibition of Chikungunya Virus Replication by 1-[(2-Methylbenzimidazol-1-yl) Methyl]-2-Oxo-Indolin-3-ylidene] Amino] Thiourea(MBZM-N-IBT) Sci Rep 2016; 6:20122. https://doi.org/10.1038/srep20122

15. Mouce BC, Poirier EZ, Passoni G., et al Interferon-Induced Spermidine-Spermine Acetyltransferase and Polyamine Depletion Restrict Zika and Chikungunya Viruses. Cell Host Microbe. 2016; 20(2):167–177 https://doi.org/10.1016/j.chom.2016.06.011

16. Mounce BC, Cesaro T, Moratorio G., et al. Inhibition of Polyamine Biosynthesis Is a Broad-Spectrum Strategy against RNA Viruses. J Virol. 2016; 90(21):9683–9692b https://doi.org/10.1128/JVI.01347-16

17. Mounce BC, Cesaro T, Vlajnic L., et al. Chikungunya Virus Overcomes Polyamine Depletion by Mutation of nsP1 and the Opal Stop Codon To Confer Enhanced Replication and Fitness. J Virol. 2017; 91(15)b https://doi.org/10.1128/JVI.00344-17

18. Mounce BC, Olsen ME, Vignuzzi M., et al. Polyamines and Their Role in Virus, Microbiol Mol Biol Rev. 2017; 81(4):1–12b. https://doi.org/10.1128/MMBR.00029-17

19. Nishimura K, Murozumi K, Shirahata A., et al. Independent roles of eIF5A and polyamines in cell proliferation. Biochem J. 2005;385(Pt 3):779–85. https://doi.org/10.1042/BJ20041477

20. Nowotarski SL, Woster PM, Casero RA. Polyamines and cancer: implications for chemotherapy and chemoprevention. Expert Rev Mol Med. 2013;15(e3): 2–21 https://doi.org/10.1017/erm.2013.3

21. Lani R. Hassandarvish P, Chiam CW., et al. Antiviral activity of silymarin against chikungunya virus. Sci Rep. 2015;5:11421

22. Lanzer W, Holowczak JA. Polyamines in vaccinia virions and polypeptides released from viral cores by acid extraction. J Virol. 1975;16(5):1254–1264.

23. Leyssen P, De Clercq E, Neyts J. Molecular strategies to inhibit the replication of RNA viruses. Antiviral Res. 2008;78(1): 9–25. https://doi.org/10.1016/j.antiviral.2008.01.004

24. Parashar D, Cherian S. Antiviral perspectives for chikungunya virus. Biomed Res Int. 2014; 631642:1–11. http://dx.doi.org/10.1155/2014/631642

25. Pegg AE. Mammalian Polyamine Metabolism and Function. IUBMB Life. 2009;61(9): 880–894. https://doi.org/10.1002/iub.230

26. Raiza NCM. Chikungunya fever: a threat to global public health. Pathog Glob Health. 2018;112(4):182–194 https://dx.doi.org/10.1080%2F20477724.2018.1478777

27. Rodríguez-Páez, L. C. La δ-N-yodoacetil-ornitina y la ε-yodoacetil-lisina como inhibidores irreversibles dirigidos al sitio activo de la ornitina descarboxilasa. An.Esc.Nac.Cienc.Biol.Mex. 1998;44(99-115).

28. Silva LA, Dermody TS. Chikungunya virus: epidemiology, replication, disease mechanisms, and prospective intervention strategies. J Clin Invest. 2017;127(3):737–749. https://doi.org/10.1172/JCI84417

29. Tuomi K, Rana A, Matyjarvi R. Synthesis of SemilIKI-Forest virus in polyamine-depleted baby-hamster kidney cells. Biochem J. 1982;206(1):115–119 https://doi.org/10.1042/bj2060113

30. Thomas T, Gallo MA, Klinge CM., et al. Polyamine-mediated conformational perturbations in DNA alter the binding of estrogen receptor to poly(dG-m^5^dC).poly (dG-m^5^dC) and a plasmid containing the estrogenesponse element, J. Steroid Biochem. Mol. Biol. 1995;54(3-4) 89–99.

31. Vargas-Ramírez AL, Medina-Enríquez MM, Cordero-Rodríguez NI., et al. N-ω-chloroacetyl-L-ornithine has in-vitro activity against cancer cell lines and in-vivo activity against ascitic and solid tumors. Anti-Cancer Drugs 2016;27(6):508–518. https://doi.org/10.1097/CAD.0000000000000353

